# Voltage-gating of mechanosensitive PIEZO channels

**DOI:** 10.1101/156489

**Authors:** Mirko Moroni, Martha Rocio Servin-Vences, Raluca Fleischer, Gary R. Lewin

## Abstract

Mechanosensitive PIEZO ion channels are evolutionarily conserved proteins whose presence is critical for normal physiology in multicellular organisms. Here we show that, in addition to mechanical stimuli, PIEZO channels are also powerfully modulated by voltage and can even switch to a purely voltage gated mode. Mutations that cause human diseases such as Xerocytosis profoundly shift voltage sensitivity of PIEZO1 channels towards the resting membrane potential and strongly promote pure voltage gating. Our data may be explained by the presence of an inactivation gate in the pore, the opening of which is promoted by outward permeation. Invertebrate (fly) and vertebrate (fish) PIEZO proteins are also voltage sensitive but voltage gating is a much more prominent feature of these older channels. We propose that the voltage sensitivity of PIEZO channels is a deep property co-opted to add a regulatory mechanism for PIEZO activation in widely different cellular contexts.

## Introduction

The mechanically-gated ion channels PIEZO1 and PIEZO2 are involved in a variety of physiological functions essential to life. Thus PIEZO1 plays a fundamental role in the development of the mouse vasculature as well as lymphatic systems (Li et al., 2014; Lukacs et al., 2015; Ranade et al., 2014a). The PIEZO2 protein is primarily found in sensory neurons of the dorsal root ganglia (DRG) and Merkel cells where its presence is essential for mechanotransduction (Coste et al., 2010; Ranade et al., 2014b; Woo et al., 2014). Mice without *Piezo2* in sensory neurons lack normal touch sensation and proprioception and severe loss of function alleles in humans are also associated with loss of proprioception and touch sensation (Chesler et al., 2016; Mahmud et al., 2017; Ranade et al., 2014b; Woo et al., 2015). Genetic ablation of either the *Piezo1* or *Piezo2* genes in mice leads to either embryonic or early post-natal lethality (Li et al., 2014; Nonomura et al., 2016; Ranade et al., 2014a). In the last few years both mouse and human genetics have revealed roles for PIEZO mechanosensing ion channels in a variety of non-sensory cellular physiology ranging from cartilage formation by chondrocytes (Lee et al., 2014; Rocio Servin-Vences et al., 2017), epithelial sheet homeostasis (Eisenhoffer et al., 2012; Gudipaty et al., 2017), growth cone guidance (Koser et al., 2016), arterial smooth muscle remodelling (Retailleau et al., 2015) as well as blood cell shape regulation (Albuisson et al., 2013).

The mouse PIEZO1 protein has been shown to be a pore forming ion channel that is directly gated by membrane stretch (Coste et al., 2012; Cox et al., 2016; Syeda et al., 2016). A high resolution picture of the mouse PIEZO1 protein was recently obtained revealing a trimeric three-bladed, propeller-shaped structure with a central pore-forming module comprising an outer helix (OH), C-terminal extracellular domain (CED), inner helix (IH) and intracellular C-terminal domain (CTD) (Ge et al., 2015). The peripheral regions of the protein are composed of the extracellular “blade” domains, 12 peripheral helices (PHs) in each subunit and intracellular “beam” and “anchor” domains. The PIEZO1 structure has facilitated biophysical exploration of the ion channel pore (Lewis and Grandl, 2015; Wu et al., 2016; Zhao et al., 2016) and experiments with chimeric structures have suggested that it is the N-terminal non-pore containing region that confers mechanosensitivity on the channel (Zhao et al., 2016). However recent experiments suggested that this subject needs further validation (Dubin et al., 2017).

Biophysical investigation of PIEZO channels have focused mostly on their activation by mechanical forces. For example, proteins like STOML3 have been identified that can dramatically increase the sensitivity of PIEZO channel to mechanical force (Poole et al., 2014; Wetzel et al., 2016). However, the role of membrane voltage in modulating PIEZO channel activity has barely been addressed despite the fact that both mammalian stretch-activated potassium channels (Brohawn et al., 2014; Dedman et al., 2009; Honore et al., 2002) as well as bacterial stretch-activated ion channels are clearly voltage sensitive (Bass et al., 2002). Here we asked whether voltage can modulate or even gate vertebrate and invertebrate PIEZO channels. We show that both PIEZO1 and PIEZO2 show significant voltage sensitivity that is dependent on critical residues in the pore lining region of the channel. We provide evidence for an inactivation gate that closes following inward permeation and under physiological conditions renders >90% of the channels unavailable for opening by mechanical force. Outward permeation of the channel is sufficient to lead to a slow conformational change that opens the inactivation gate. Pathological human mutations in *Piezo1* primarily weaken the inactivation gate and render channels insensitive to voltage modulation. The same mutations also allow PIEZO1 to behave as a voltage-gated ion channel in the absence of mechanical stimuli. Finally we show that the biophysical properties of both invertebrate (*D. melanogaster*) and other vertebrate (*D. rerio*) PIEZOs are much more reminiscent of classical voltage-gated channels than mammalian PIEZOs.

## RESULTS

### PIEZO1 open probability is increased at positive voltages

PIEZO1 channels classically show a linear I/V relationship, but current inactivation is much slower at positive holding potentials compared to negative voltages (Coste et al., 2010; Zhao et al., 2016). Here we subjected excised outside-out patches from N2A cells containing over-expressed mouse PIEZO1 to an I/V protocol from -100mV to +100mV in symmetrical Na^+^, without divalent cations, while stimulating the patch with a saturating pressure pulse using a fast pressure clamp system. Under these conditions PIEZO1 has been shown to display a linear I/V relationship measured between -50mV to +50mV (Coste et al., 2010; Zhao et al., 2016). However, at more positive voltages the I/V relationship showed strong outward rectification (rectification index I_-60mV_/I_60mV_ 5.3 ± 0.6, 12 cells) (Figure 1A). One explanation for this behavior might be that single PIEZO1 channels conduct more ions outward than inward, a phenomenon seen in other channels such as Glycine and GABAA channels (Moroni et al., 2011). However, single channel measurements under identical conditions revealed that PIEZO1 channel outward conductance was actually smaller than the inward conductance (Figure 1B). Thus, changes in single channel conductance cannot account for the novel outward rectification seen starting at holding potentials >0 mV (Figure 1B).

**Figure 1.**
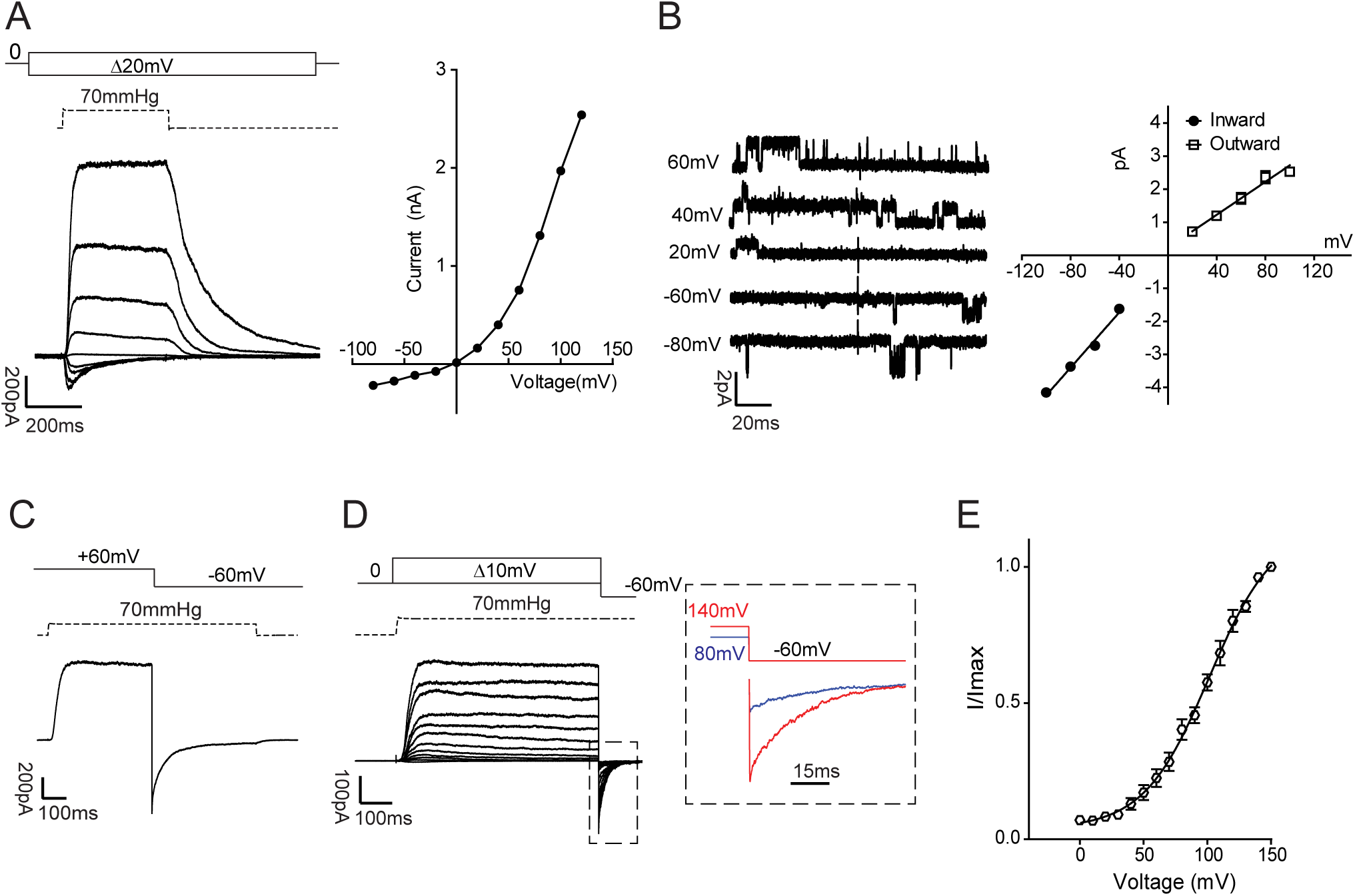
Rectification and voltage modulation of pressure-mediated PIEZO1 currents. A) *Left*. Example traces of currents elicited at a constant saturating pressure (70mmHg) and at increasing voltages (in 20mV steps from -100mV to 100mV) in symmetrical Na^+^ from excised (outside-out) patches overexpressing mPIEZO1 in N2a cells. *Right.* Peak currents are plotted against voltage to show an I/V relationship. Note the outwardly rectifying behaviour. B) *Left.* single channel openings were recorded at negative and positive voltages to obtain slope conductance values. *Right.* Linear regressions from individual patches were averaged and pooled. Inward slope conductance was significantly higher than outward slope conductance (41.3 ± 0.9 pS and 27.1 ± 1.2 pS respectively, 3 cells. Student’s *t* test *P*= 0.0006). C) Example trace of instantaneous currents recorded upon switching voltage from +60mV to -60mV during the application of a 70mmHg pressure stimulus (rectification index 1.13 ± 0.06 (n=8)) Capacitance currents were digitally subtracted. D) Current responses to a pressure stimulation of 70mmHg during 300ms voltage steps ranging from 0 to 150mV, followed by a repolarization step to -60mV to obtain tail currents. The inset shows an expanded example of tail currents at -60mV originated from a pre-stimulation step at 80 (blue) and 140mV (red) in presence of 70mmHg of pressure. E) Tail currents from individual cells were normalized to their maximum and fitted to Boltzamann relationship (V_50_ = 91.9 ± 3.2 mV, slope 22.2 ± 0.9, 12 cells). Pooled data are shown as mean ± SEM.

If the apparent outward rectification behavior of PIEZO1 channels were due to intrinsic pore properties then measurements of instantaneous currents upon voltage changes should give rise to the same rectification index. We took advantage of the slow inactivation kinetic of PIEZO1 at positive voltages to trigger channel opening at +60mV with a saturating pressure pulse (70mmHg). Upon reaching steady-state activation the voltage was then stepped to -60mV and the peak instantaneous current measured. Using this protocol we found that the rectification index (I_ins-60mV_/I_60mV_) was 1.13 ± 0.06 (n=8) (Figure 1C), suggesting that the pore region can conduct macroscopic current approximately equally in both directions and that the ion permeation pathway cannot account alone for the rectification of PIEZO1. We speculated that not only pressure but also voltage could affect the apparent open probability of PIEZO1. We employed tail current protocols to understand whether voltage directly affects apparent channel open probability. A saturating pressure pulse was applied simultaneously with voltage steps ranging from 0 mV to +150 mV (Figure 1D). Upon reaching steady state activation, the voltage was stepped to -60mV in the continuous presence of pressure. We found that the amplitude of inward (tail) currents upon repolarization to -60mV increased proportionally to the magnitude of the positive pre-pulse voltage, indicating a voltage-dependent increase in apparent open probability. A Boltzmann fit of this data (Figure 1E) established that voltage is a major contributor to the gating of PIEZO1 and that depolarized voltages increase open probability (V_50_ = 91.9 ± 3.2 mV, slope 22.2 ± 0.9, 12 cells). Careful examination of the Boltzmann fit indicates that at physiological resting membrane potentials (<0 mV) the apparent open probability or PIEZO1 channel availability, is only a tiny fraction (<10%) of the maximum that can be made available by a depolarizing pre-pulse (Figure 1E).

#### Outward permeation resets PIEZO1 kinetics

Mechanical stimulation of PIEZO1 via pressure, cell poking or substrate deflection at negative potentials shows fast inactivation (Coste et al., 2010, 2012; Poole et al., 2014; Rocio Servin-Vences et al., 2017; Zhao et al., 2016). Using pressure clamp we also measured fast inactivation time constants (τ_inact_ = 60.9 ± 3 ms, 8 cells). It has also been reported that repetitive mechanical stimulation can drive PIEZO1 channels into an irreversible non-inactivating and desensitized state (Gottlieb et al., 2012; Poole et al., 2014). Recordings of PIEZO1 macroscopic currents shows that this desensitized state, induced by repetitive stimulation (every 1 s), is characterized by a loss of inactivation as reflected in a reduction in the ratio of peak to steady-state current (Figure 2A). We next asked whether positive holding potentials affect the behavior of PIEZO1 currents to repetitive stimuli. Interestingly, mechanical stimuli applied every second at +60mV induced no loss of PIEZO1 inactivation and only minimal desensitization (Figure 2B,D). PIEZO1 is involved in a variety of biological processes where pressure, stretch, or shear stress are constantly monitored (Li et al., 2014; Ranade et al., 2014a; Retailleau et al., 2015) and we reasoned that there must be a mechanism to release PIEZO1 channels from a desensitized and non-inactivating state. By analogy to fast inactivating voltage-gated sodium channels where a long lasting hyperpolarization allows channels to rapidly exit inactivation, we reasoned that positive voltages could reset PIEZO1 channels. We thus applied constant pressure pulses at alternating negative and positive voltages (Figure 2C). Desensitization of PIEZO1 currents induced by repetitive stimuli was completely abolished as the current peak recorded at -60mV remained constant over many stimuli (Figure 2C,D). Positive voltages also prevented the channel from acquiring non inactivating kinetics (Figure 2C). Since different channel states likely correspond to distinct structural channel conformations it is plausible to assume that positive voltage together with outward permeation may hold the channel in a fully active conformation preventing the desensitized state.

**Figure 2.**
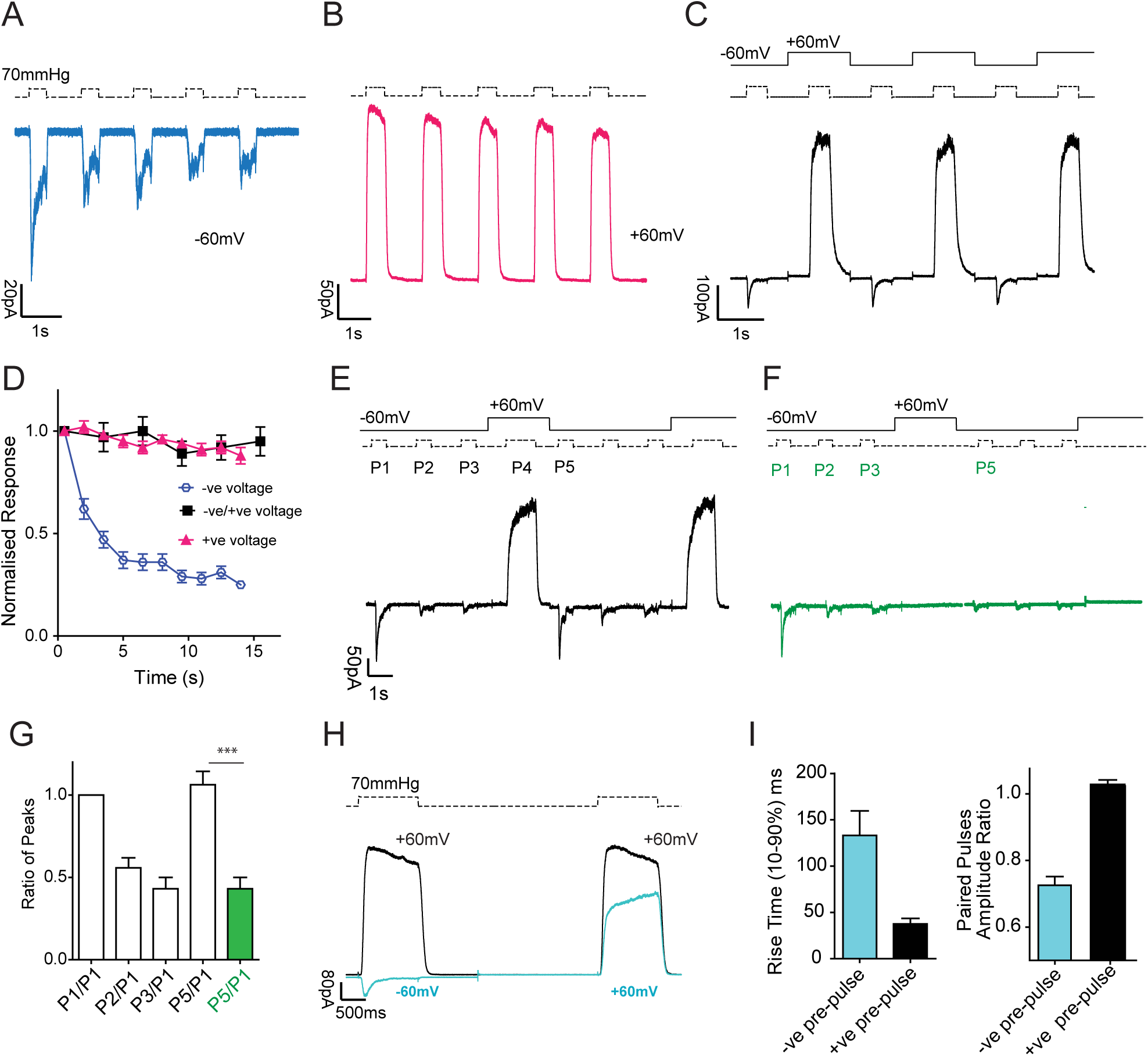
Inactivation and desensitization of PIEZO1 can be reset by outward permeation. Repetitive pressure stimulations desensitizes PIEZO1 at -60mV (A) but not at +60mV (B). At -60mV the fifth stimulus is much smaller than the first one and the current shows a typically transformation from an inactivating to a non-inactivating behavior (decreased peak current and low peak/steady state current ratio). C) Alternating pressure pulses at -60mV and +60mV abolishes desensitization and prevents PIEZO1 from entering a non-inactivating state. D) Peak currents in A (negative pulses), B (positive pulses) and C (negative pulses) were normalized to the first response and plotted against time. Note the lack of desensitization for pulses during pressure stimulation at alternating voltages. E) 3 pressure pulses (P1, P2, P3) at -60mV were applied to patches expressing PIEZO1 to drive the channel into an inactive and desensitized state, followed by 1 positive pressure pulses (P4) at +60mV. The sequence was repeated twice. Outward permeation (P4) recovers PIEZO1 initial current (P1) and resets its time course for inactivation, as shown by the ratio between the current amplitude of P5/P1, shown in G). F) The stimulation sequence in E was repeated in absence of pressure to prevent channel opening and permeation when switching to a positive voltage (P4). The recovery from desensitization does not occur in absence of a permeation event, underlining the importance of outward permeation for resetting channel kinetics. G) The pulses ratio P1/P5 for G (white) and F (green) are shown and are statistically significant *P*<0.0001, n=10. H) Paired pulses protocol of a pressure stimulus at +60mV preceded by either a pressure step at -60mV (blue trace) or at +60mV (black trace). Note how the direction of the first current stimulus affects the amplitude and the time course of activation of an identical second step at +60mV. I) The rise time and the amplitude ratio of the second stimulus at +60mV preceded by a pulse at either negative (light blue) or at positive voltage (black) are compared. Both rise time and current amplitude ratio are significantly different (Students t-test *P*<0.001). The slow rise time of the current at +60mV most likely underlies a slow conformational change and its exiting from an desensitized state.

We next asked whether outward currents could restore an active conformational state after the channel had been forced into a desensitized state. To test this hypothesis we applied three pressure pulses at negative voltages followed by one pressure pulse at a positive voltage. The whole sequence was repeated twice without a pause (Figure 2E). After three pressure pulses (P1-P3) PIEZO1 currents showed marked desensitization, measured as decreased current amplitude compared to the first pulse P1 (Figure 2E). Once outward permeation of PIEZO1 occurs by stepping to +60 mV the current amplitude of pulse 5 (P5) measured at -60 mV was identical in amplitude and kinetics to P1 (Figure 2E and G) (P1 τ_inact_ 62.3 ± 2.1 ms, P5 61.4 ± 3.1 ms 8 cells). This experiment nicely demonstrates how voltage or outward permeation can reset the channel to its fully active state. We next asked whether it is permeation or positive voltage that is necessary to reset desensitization. We repeated the experiments as above, but did not apply pressure pulse P4 while holding the patch at +60 mV. Under these conditions, no outward current was measured and PIEZO1 channels did not recover from their desensitized state; the ratio of P5/P1 was 0.25 ± 0.5 (n=10) (Figure 2F and G). These experiments established that the loss of inactivation is not an irreversible event, as has previously been suggested (Gottlieb and Sachs, 2012), and that outward ion conduction is sufficient to reset normal PIEZO1 channel properties. We noticed that the transition between the desensitized state (after 3 pressure pulses at -60mV, P1-P3 Figure 2E) and a “recovered” active state (P5) (after the pulse at +60mV, P4 Figure 2E) may require a slow conformational change as reflected in the slow rise in current amplitude during P4. To explore this idea we made use of a paired pulse protocol. We compared peak currents elicited by pressure pulses at +60mV that were preceded by a pressure pulse at -60 mV (blue) or +60 mV (black) (Figure 2H). The currents evoked by the second pressure pulse showed a 3-fold slower rise time and 25% reduced amplitude when the preceding pressure pulse was delivered at -60 mV compared to +60 mV (Figure 2H and I). Together these observations demonstrate that transitions between active and inactive conformational states for PIEZO1 are reversible. More striking was our finding that resetting of inactivation and removing desensitization only requires outward ion permeation through the PIEZO1 channel.

#### Removal of inactivation requires outward permeation

Since outward permeation resets channel properties we hypothesized that an outward movement of ions could remove inactivation in a manner proportional to the driving force. To investigate how outward currents remove inactivation we recorded currents elicited by pressure pulses at -60mV that were preceded by conditioning pressure steps with increasing driving force (voltages from +20 to 140mV). If our hypothesis is correct the magnitude of the second pressure-induced current measured at -60mV should be dependent on the magnitude of current evoked by the preceding conditioning step. Indeed we found that the larger the outward current measured in the conditioning step the larger the subsequent inward current measured at -60 mV (Figure 3A,B). By plotting the conditioning voltage versus the peak current amplitude at -60mV we calculated that removal of inactivation had a *V*_50_ of 85.5 ± 5 mV (n=10). Thus the inactivation state of PIEZO1 can be efficiently reversed and the transition between the inactive and the active state occurs in a manner proportional to the amount of ions that flow through the channel. This provides an important mechanistic link between permeation and inactivation and implies the presence of an inactivation gate along the permeation path. The opening of this putative inactivation gate would depend on the amount of ions that flow outward through the pore. Furthermore, this inactivation gate can be forced open by increasing the outward driving force during permeation, thus establishing an essential role for permeating ions in the gating of PIEZO1 (Figure 3E).

**Figure 3.**
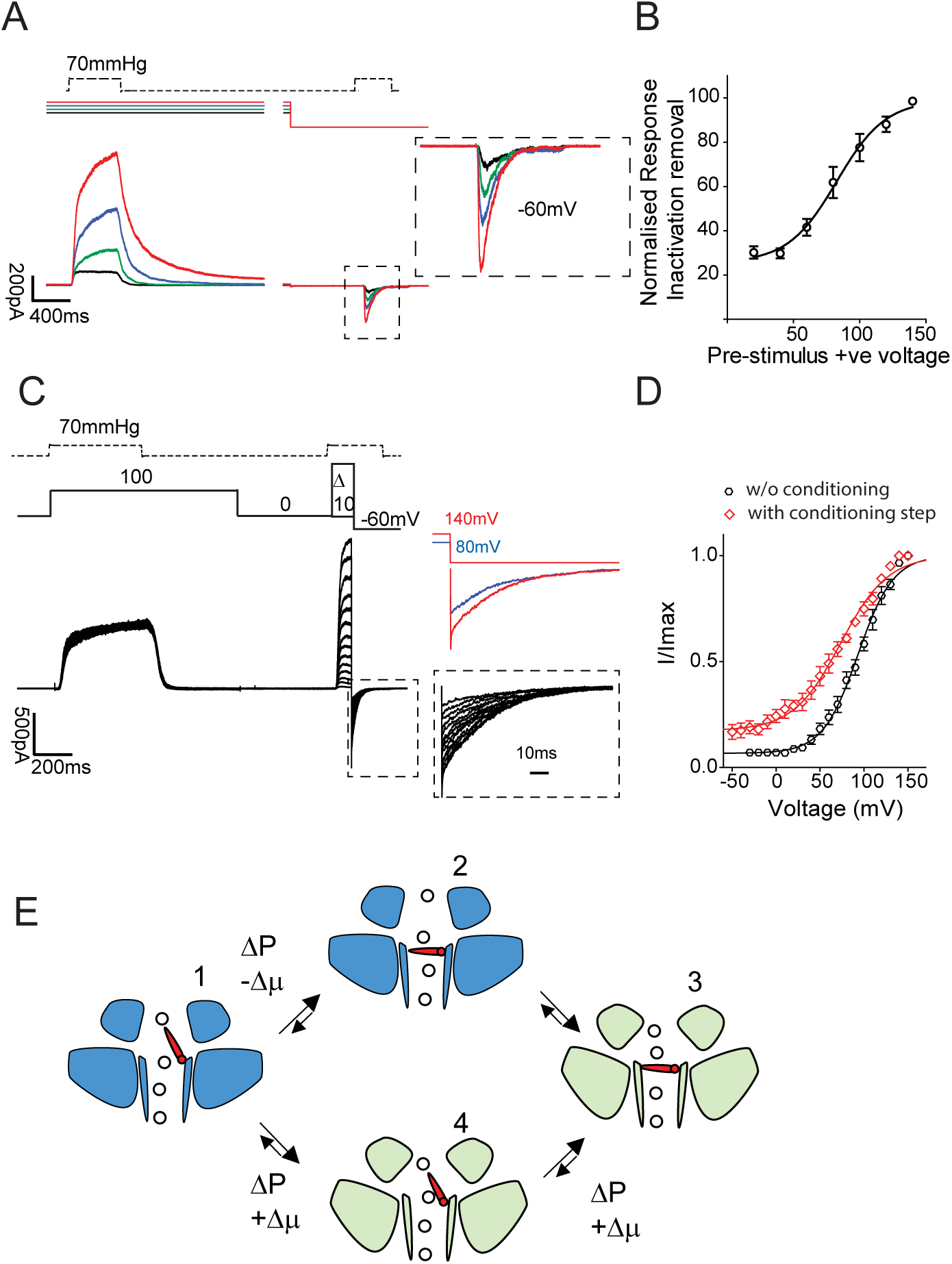
Increasing outward permeation determines the number of channels available for activation. A) A constant pressure pulse at -60mV was preceded by a family of constant pressure stimulation (70mmHg) at increasing voltages (20mV black, 40mV green, 60mV blue, 80mV red). The current amplitude of the second pressure stimulation depends on the driving force applied during the preceding step, showing that the larger the applied driving force the greater the relief from inactivation. B) The conditioning stimulus voltage was plotted against the normalized current amplitude of the currents recorded at -60mV. Single cells were fitted individually to a Boltzmann fit. The data shown represent pooled data from 10 cells (*V*_50_ = 85.5 ± 5 mV). C) Tail current protocol as in Fig 1D preceded by a conditioning pressure pulse at +100mV to remove ~80% of inactivation. D) Tail currents from individual cells were normalized to their maximum and fitted to a Boltzmann relationship (V_50_ = 90.6 ± 2.9 mv, slope 24.6 ± 1.6, n=12 cells without conditioning step, black trace, V_50_ = 68.3 ± 10.2 mV, slope 43.7 ± 2.3, 5 cells, with conditioning step at +100mV, red trace, Student’s *t*-test *p*=0.0036). Pooled data are shown as mean ± SEM. E) Proposed transition between active (blue) and inactive state (light green of PIEZO1). Activation by pressure (ΔP) and negative electrochemical gradient (-Δ*μ*) drive the channel into an inactive state (light green). Pressure and positive electrochemical gradient (+Δ*μ*) reset the channel to an active state by opening an inactivation gate. The transition 3-4-1 is a slow conformational change as suggested by the rise time and amplitude in 2 H and I.

We extended our findings using tail current protocols. We repeated the tail current protocol introduced in Figure 1F but added a conditioning pressure pulse at +100mV. The conditioning pressure pulse at positive potentials removes ~80% of the inactivation (Figure 3C), and by plotting tail current amplitudes against voltage we measured a substantial leftward shift in the current-voltage relationship of ~20mV (90.6 ± 2.9 mv, n=12 without conditioning step, black trace, and 68.3 ± 3.7 mV n=5, with conditioning step, red trace, Student’s *t*-test *p*=0.0036). Thus, outward permeation of the channel not only removes inactivation but can also increase the apparent open probability of the channels. The shift in I/V relation induced by a positive pre-pulse effectively leads to a three-fold increase in the number of available channels at physiological membrane potentials (< 0 mV) from 5 to 15% (Figure 3D).

#### Altered voltage modulation of PIEZO1 in human disease

Missense mutations in the coding sequence of Piezo1 lead to diseases such as Xerocytosis and lymphatic dysplasia (Albuisson et al., 2013; Andolfo et al., 2013; Fotiou et al., 2015; Lukacs et al., 2015; Shmukler et al., 2014; Zarychanski et al., 2012). Mutations causing Xerocytosis lead to a slowing of channel inactivation to membrane stretch or in response to cell poking (Albuisson et al., 2013; Lukacs et al., 2015). We introduced the missense mutation R to H at position 2482 in the mouse cDNA (human R2456H), a residue that is located at the bottom of the putative pore forming inner helix in the C-terminal domain of mPIEZO1 (Ge et al., 2015; Zhao et al., 2016). The human R2456H mutation was shown to display the strongest slowing of inactivation phenotype amongst the known Xerocytosis mutants (Albuisson et al., 2013), and we observed a similar slowing of inactivation for mouse PIEZO1 with the orthologous R2482H mutation (Figure S1A). We examined desensitization properties of this mutant protein and measured a decrease in the peak current amplitude upon repetitive mechanical stimuli at negative (-60mV) exactly as seen in the wild type (Figure S1B). Current amplitude was also maintained when stimulating at positive holding potentials (+60mV) and alternating positive/negative voltages (-60mV/+60mV), exactly as seen for the wild type channel (Figure S1C,D). The slow transition out of the inactive/desensitized state (measured from the rise time of a pressure pulse at +60mV after 3 pulses at -60mV) was also similar to the wild type (Figure S1E,F).

We next used tail current protocols to measure the apparent open probability of the channels and observed that the R2482H mutation was associated with a large 50 mV leftward shift in the current-voltage relationship compared to wild type. When R2482 was mutated to a lysine (K), the current-voltage relationship was shifted even further to negative voltages (R2482H: *V*_50_ 42.0 ± 3.7 mV, 19 cells, R2482K: *V*_50_ 38.0 ± 2.7 mV, 6 cells, both mutants were significantly different from the wild type *V*_50_ 91.8 ± 3.2 mV, 7 cells, *p*<0.0001, one-way ANOVA and Dunnett’s posttest) (Figure 4). Interestingly, both the R2482H and R2482K mutants showed a much higher apparent open probability at negative voltages than the wild type channels (compare tail currents in Figure 4 A, and also the bottom of the current-voltage relationship in Figure 4B). We next asked whether other Xerocytosis disease mutations in the *Piezo1* gene can also alter PIEZO1 voltage sensitivity. We thus introduced the following mutations in the mouse cDNA: R1353P (human R1358P (Albuisson et al., 2013)) is located quite distant to the pore on a predicted helix not resolved in the structure (Ge et al., 2015), and A2036T (human A2020T (Albuisson et al., 2013)) is located on a peripheral membrane helix that is resolved in the structure and finally T2143M (human T2127M located in the middle of the anchor domain (Figure 4C). In contrast to the R2482H and R2482K Xerocytosis mutants, the peripherally located A2036T and R1353T mutations had no effect on the voltage sensitivity of the channel when compared to wild type (Figure 4A,B). Despite the lack of effect on voltage sensitivity both these mutations did significantly slow the inactivation kinetics of the channel (Figure 4D). In contrast, the T2143M anchor site mutation had a very similar effect to the R2482H mutation in dramatically altering the voltage sensitivity of the channel. Structurally, the mutated threonine (2143) is predicted to be located just 17 Å distant from the arginine (24) located on the adjacent subunit, suggesting that both residues participate in spatially restricted domain of the channel that control voltage dependent gating of PIEZO1 channels. Thus the Xerocytosis disease mutations allowed us to identify distinct parts of the channel that dramatically influence voltage-dependent gating distinct from channel inactivation kinetics.

**Figure 4.**
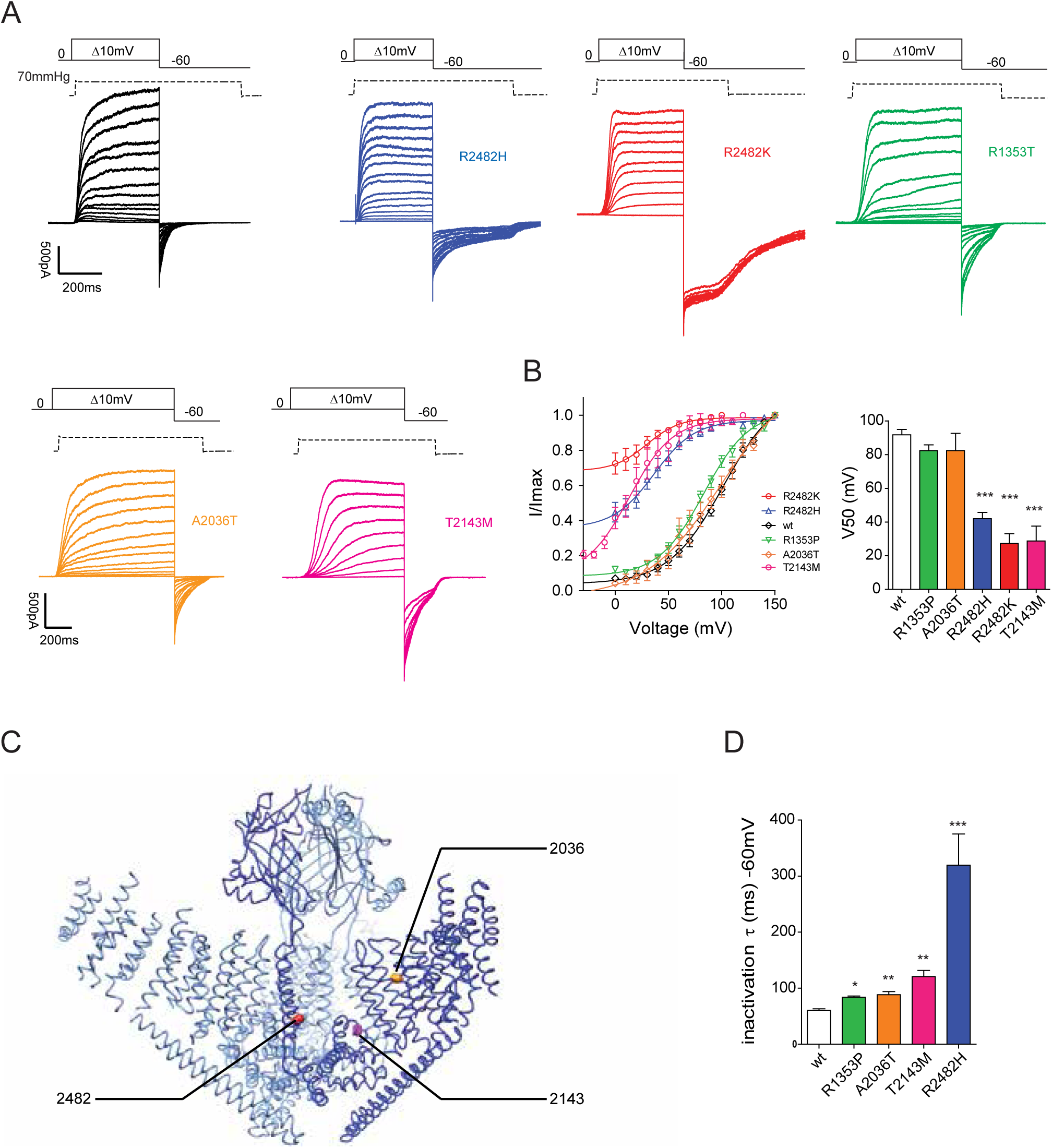
Mutations causing a Xerocytosis in humans alter voltage modulation of PIEZO1. A) Current responses to a pressure stimulation of 70mmHg during 300ms voltage steps ranging from 0 to 150mV, followed by a repolarization step to -60mV to obtain tail currents for wild type PIEZO1 (black), R2482H (blue), R2482K (red), R1353T (green), A2036T (orange), T2143 (magenta) mutants. The R2482K mutant shows almost no voltage modulation. B) Tail currents from individual cells were normalized to their maximum and fitted to a Boltzmann relationship (wt: *V*_50_ = 91.9 ± 3.2 mV, slope 22.2 ± 0.9, 12 cells; R2482H *V*_50_ = 42.0 ± 3.7 mV, slope 14.4 ± 1.5, 19 cells;; R2482K *V*_50_ 27.2 ± 5.9 mV, slope 12.2 ± 1.5, 10 cells, R1353P *V*_50_ = 82.4 ± 8.8 mV, slope 20.1 ± 1.3, 8 cells, A2036T *V*_50_ = 82.4 ± 10.2 mV, slope 22.4 ± 0.9, 5 cells, T2143M *V*_50_ = 28.7 ± 8.8 mV, slope 17.6 ± 2.0, 8 cells). Pooled data are shown as mean ± SEM.C) 3D structure of the trimeric mPIEZO1 showing the position of the mutants involved in Xerocytosis. R1353 is not present in the available structure and it is not shown. R2482 is in the bottom of the inner helix, A2036 is located in a peripheral helix, T2143 is in the anchor domain. D) Inactivation time constant of wild type mPIEZO1 and of the Xerocytosis mutants in response to a saturating pressure pulse at -60mV. All mutants showed a significantly different inactivation time course from the wild type channel. (See also Figure S1)

#### Voltage modulation of PIEZO2

PIEZO2 has a fundamental role in sensory neuron mechanotransduction where it is necessary for mechanoreceptor function and the sense of light touch and proprioception in humans and mice (Chesler et al., 2016; Coste et al., 2010; Maksimovic et al., 2014; Ranade et al., 2014b; Woo et al., 2015). Mechanically-activated PIEZO2 currents inactivate faster than PIEZO1 currents (Coste et al., 2010; Poole et al., 2014). We next asked whether the apparent open probability of PIEZO2 channels are modulated by voltage in a similar way to PIEZO1. However, in excised or cell attached-patches from cells overexpressing PIEZO2 we were unable to record stretch-activated currents (Figure 5C). A similar observation has been reported for endogenously expressed PIEZO2 in Merkel cells (Ikeda and Gu, 2014). This suggests that PIEZO2 might not be able to respond directly to membrane stretch or that PIEZO2 might need to be associated with other molecules in the membrane or at the membrane substrate interface to be gated by mechanical stimuli, a phenomenon described for TRPV4 (Rocio Servin-Vences et al., 2017). Using pressure to gate channels in excised patches has many experimental advantages due to small patch capacitance, no space clamp issues, and the ability to control both intracellular and extracellular ion concentrations. We show that voltage-dependent modulation of PIEZO1 depends on the permeation pathway and is strongly affected by the mutation of residues around the pore area in the C-terminal region (K2188—E2547) (Zhao et al., 2016). We decided to generate a stretch-activated chimeric PIEZO1/PIEZO2 protein (P1/P2 chimera) by fusing the N-terminal sequence of PIEZO1 up to and including the so called anchor region (aa 1-2190) with the remaining downstream sequences of PIEZO2 (aa Y2472-N2822). The C-terminal PIEZO2 sequences of the chimera include the last two transmembrane spanning domains that are thought to form the channel pore, the so called inner helix (IH) and outer helix (OH) as well as the C-terminal extracellular domain (CED) and C-terminal domain (CTD) (Figure 5A). To study the chimeric channel under the same controlled experimental system we used a CRISPR/Cas9 approach to generate a N2a cells line in which the mouse *Piezo1* gene had been knocked out (referred to as N2a *^Piezo1^*^-/-^ cells, see Methods and Figure S2). We overexpressed the P1/P2 chimeric channel and could in using whole cell patch clamp record robust currents to soma-indentation that had similar amplitudes to wild type PIEZO2 currents (Figure 5A,B). More importantly, in excised patches from cells overexpressing the P1/P2 chimeric channel we could record robust stretch-activated currents (Figure 5C). The pressure sensitivity of the P1/P2 chimeric channel measured by applying series of fast pressure clamp pulses was very similar to that of PIEZO1 channels, thus from pressure-current plots we calculated a *P*_50_ of 37.6 ± 4.6 mmHg (10 cells) for the P1/P2 chimera compared to 38.9 ± 3.1 mmHg, 21 cells for PIEZO1 (Figure 5C,D). Interestingly, the inactivation kinetics of currents measured from P1/P2 chimeric channels were ~ 3 fold faster than PIEZO1 (26.3 ± 6.2 ms, 4 cells for P1/P2 channel and 60.9 ± 3.0 ms, 9 cells for wt PIEZO1) (Figure 5B, E), and thereby showed similarities to the faster inactivation kinetics PIEZO2 channels activated by soma indentation. This experiment indicates that the domains that confer membrane-stretch sensitivity to PIEZO1 reside within the N-terminal domain of the protein. Most importantly, we found that the last two transmembrane domains are crucial for setting the speed of inactivation. Strikingly, also the chimeric P1/P2 channel showed an apparent rectifying behavior when patches were subject to an I/V protocol as shown for PIEZO1 (Figure S3A), suggesting that also PIEZO2 might be controlled by voltage in a similar way to PIEZO1. To test our hypotheses we repeated the experiments in Figure 2 on the P1/P2 chimera. As expected the desensitization of PIEZO2 also occurred at negative voltages but it hardly occurred at positive potentials even after 10 repetitive stimulations at 1Hz (Figure S3B). As for PIEZO1 the P1/P2 chimera passed the same amplitude of current when stimulated at alternating negative and positive voltages. We further asked whether outward permeation along the PIEZO2 pore would also drive the channel out of desensitization by opening of a putative inactivation gate. Indeed PIEZO2 current amplitude can be restored to its initial value by outward permeation as shown by a protocol in which a series of pressure pulses are delivered at positive or negative voltage (Figure S3C).

**Figure 5.**
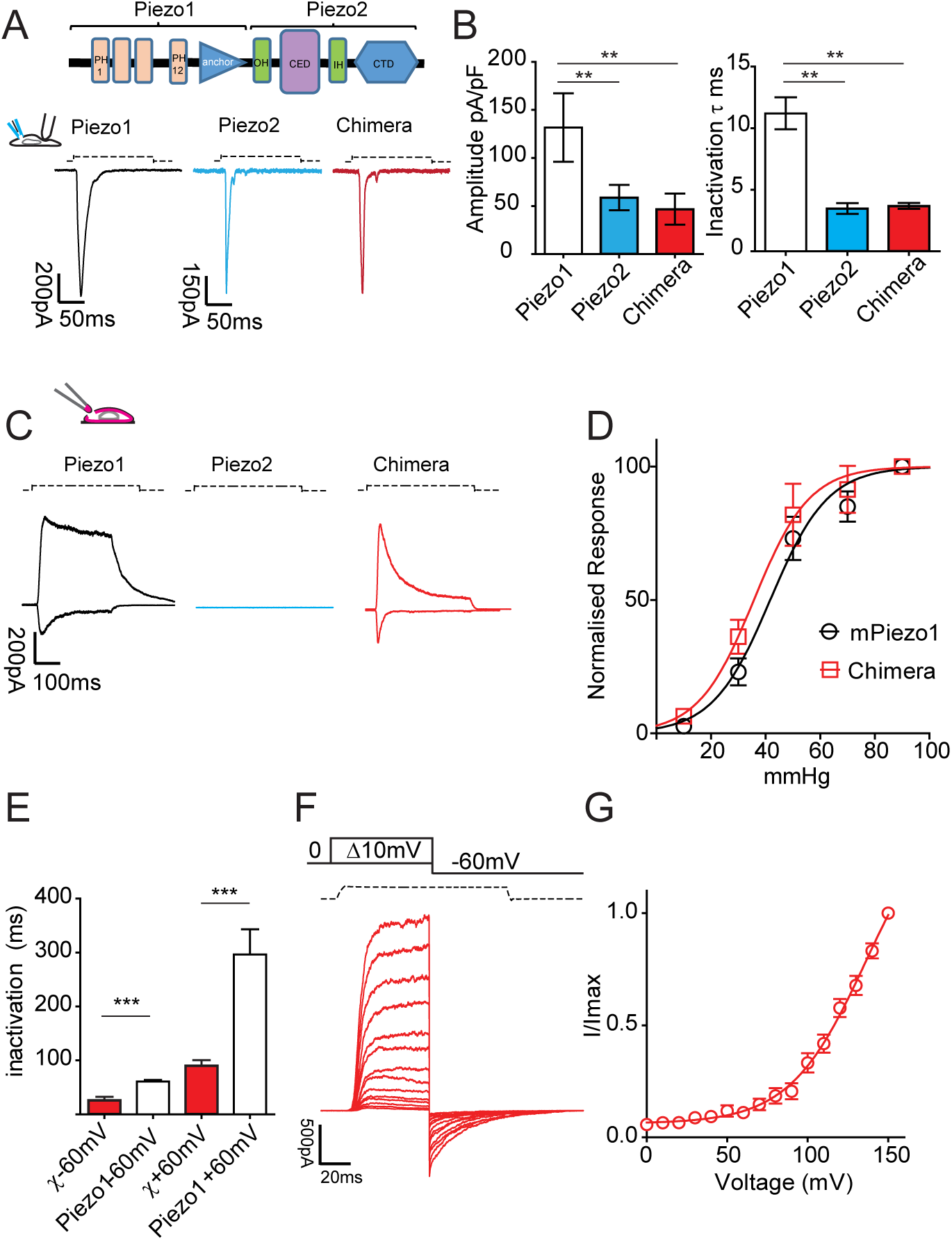
PIEZO1/PIEZO2 chimera responds to membrane stretch and it is modulated by voltage. A) A chimeric PIEZO1-PIEZO2 receptor was constructed by fusing the N terminal region of PIEZO1 and the last two transmembrane domains of PIEZO2 including the CED (C terminal extracellular domain) region. PIEZO1, PIEZO2 and the chimeric channel were overexpressed in N2a *^Piezo1^*^-/-^ cells (See also Figure S2). Cells were clamped at -60mV and subjected to soma indentation. B) The maximum current amplitude (normalized for the capacitance of each cell) is plotted. The time course of inactivation for each construct was fitted to a mono exponential function. PIEZO1 inactivation was ~3 fold slower than PIEZO2 and the chimeric receptor (ANOVA, Dunnett’s post-test P<0.05). The chimeric receptor showed a current amplitude and a time course of inactivation identical to PIEZO2. C) Typical response of the chimeric channel and wild type PIEZO1 to membrane stretch in outside-out patches. Outside-out patches pulled from cells overexpressing PIEZO2 did not respond to stretch stimulation. D) Pressure-response relationship of the chimeric receptor is not different from PIEZO1 (chimera 38.9 ± 4.6 mmHg, 10 cells, PIEZO1 38.9 ± 3.1 mmHg, Student’s *t*-test *P*=0.78). E) The decay of inactivation for pressure-mediated responses at +60mV and -60mV is plotted for the chimeric and PIEZO1 channels. Also for pressure-mediated responses the kinetic of inactivation of PIEZO1 remained ~3 fold slower than the chimeric channel F) Current responses to a pressure stimulation of 70mmHg during 60ms voltage steps ranging from 0 to 150mV, followed by a repolarization step to -60mV to obtain tail currents. G) Tail currents from individual cells were normalized to their maximum and plotted against voltage. Pooled data are shown. (See also Figure S3)

Last, we asked whether also the P1/P2 chimera could be modulated by voltage in a manner similar to wild type PIEZO1. Figure 5F and G show that as for PIEZO1, the chimeric channel could also produce robust tail currents at -60mV. The amplitude of the tail currents increased with the preceding step and the apparent open probability increased up to voltages as high as 150mV. These data show that the pore region of PIEZO2, like PIEZO1, is also capable of sensing changes in voltage.

#### Direct voltage gating of PIEZO1

We established that the apparent open probability of PIEZO1 and PIEZO2 can be modulated by voltage so that depolarization dramatically increased the number of membrane stretch-activated channels. Normally, inward ion permeation drives the channels into an inactive state that can be removed by outward ion conduction, perhaps by opening an inactivation gate. We hypothesized that a prerequisite for voltage alone to gate PIEZO1 would be a fully open inactivation gate. The R2482H/K mutant channels might fulfill such a condition as we had observed a very high apparent open probability at voltages where wild type channels are mostly not available (Figure 4). We mechanically-activated PIEZO1 channels during step depolarizations from 0 to +80 mV and as expected outward currents start to decay immediately after cessation of the pressure pulse reflecting deactivation of the channels which was slower at more depolarized potentials (Figure 6 and (Coste et al., 2010)). However, the R2482H and R2482K mutant channels showed an unexpected additional feature: current deactivation was seen immediately after the pulse, but 20-40 ms later the current started to reactivate as evidenced by sag and increased or maintained outward current (Figure 6A). The reactivation of the mutant PIEZO1 outward current was only seen at more depolarized potentials and we estimated the mean threshold for this effect to be 34.4 ± 3.4 mV (9 cells). This suggests that voltage might play a role in directly (re-)opening PIEZO1 channels, as it normally occurs in voltage gated ion channels. We hypothesized that during the sag phase outward permeation leads to a structural change in the mutant PIEZO1 channel, which promotes a new stable conformation in which the inactivation gate is fully open and in which the channel might respond directly to voltage without the need of a pressure stimulus. Polymodality in activation has been previously described for ion channels. Therefore, we hypothesized that PIEZO1 could sequentially switch between two modes of activation: a pressure-gated and a voltage gated mode.

**Figure 6.**
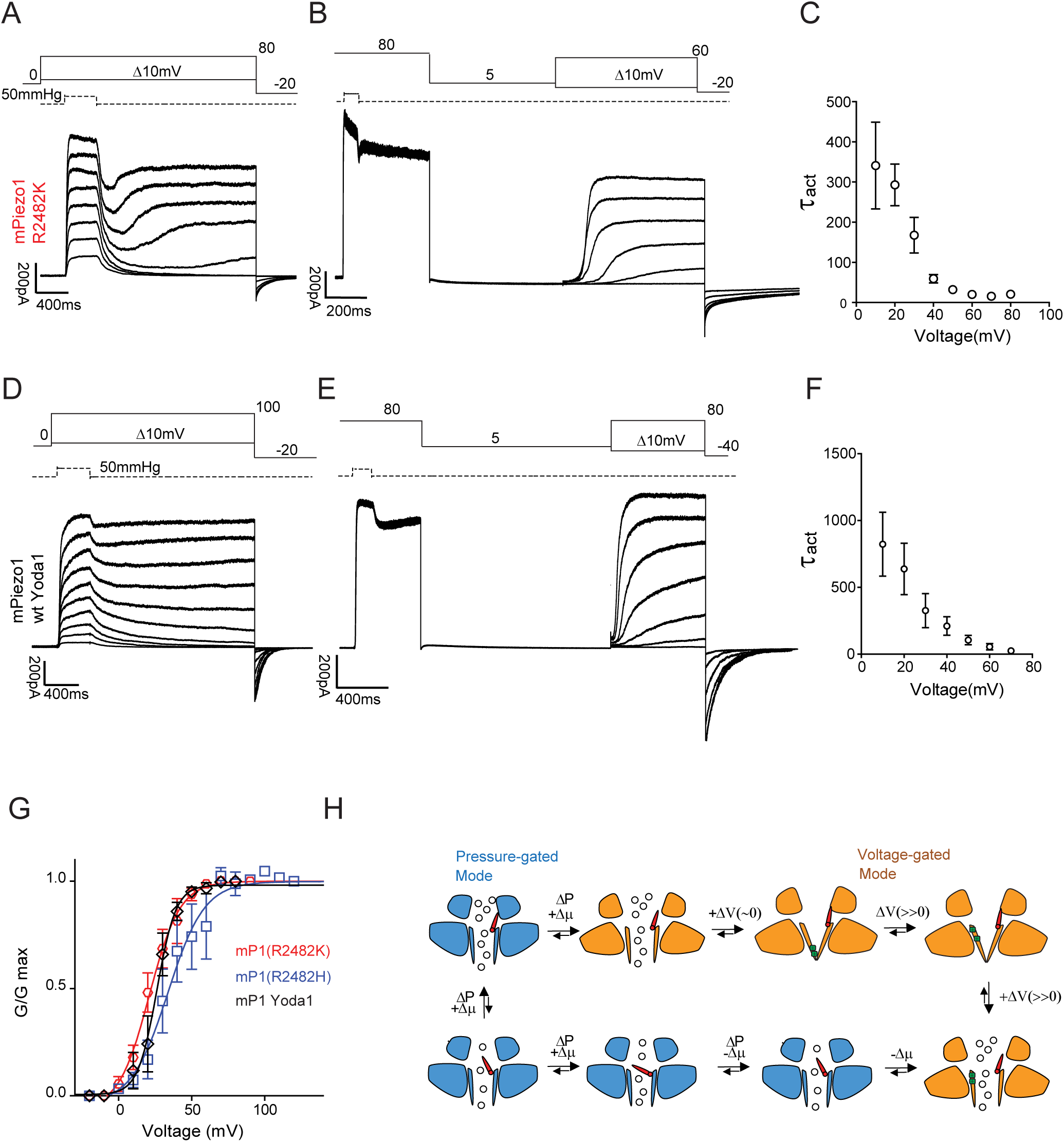
PIEZO1 transitions between a pressure-gated and a voltage-gated mode. A) The PIEZO1 R2482K mutant was subject to a family of pressure stimulations at increasing voltages (from 10 to 80mV) and the deactivation phase was extended to 2 seconds. After an initial deactivation PIEZO1, at voltages above 40mV, undergoes a reactivation phase. Reactivation time constants are plotted in Supplementary Fig 3D. B) A saturating pressure pulse at 80mV was applied to force the channel to reactivate and switch to a voltage-gated mode. Following a 2 s deactivation period at 5mV (to prevent the inactivation gate from closing) a family of 1 s steps at increasing voltages (in absence of pressure) was applied, showing that PIEZO1 R2482H/K can be activated by the sole application of voltage as in the case of canonical voltage-gated ion channels. C) The time course of activation increased at more depolarized voltages. The time constants of activation for mutant R2482K are plotted against the voltage. D) wild type PIEZO1 undergoes reactivation in the presence of a low concentration of the gating modifier Yoda1 (5μM) when included in the intracellular solution. The same voltage/pressure protocol was applied as in A. E) PIEZO1 can be activated by voltage in presence of Yoda1 (5μM) intracellularly in absence of any pressure stimulation. The same voltage/pressure protocol was applied as in B. F) The time course of activation decreased at more depolarized voltages as it occurs in voltage-gated ion channels. G) Conductance-voltage relationships for PIEZO1 R2482K (red), R2482H (blue), wt + Yoda1 5μM (black) in voltage-gated mode were fitted to a Boltzmann equation. H) Proposed model for gating transitions of PIEZO1. Starting from the bottom left: PIEZO1 inactivation gate opens during outward permeation (+Δμ) and application of pressure (ΔP) (red inactivation gate partially tilted upward). A further and persistent depolarization (>40mV for R2482 mutants and PIEZO1 in presence of Yoda1) can overcome inactivation and induce a reactivation of the channel (inactivation gate completely open) and a switch to a voltage-gated mode (in orange). Deactivation of the channel at voltages close to 0 lead the channel to close. Further depolarization causes a movement of gating charges and opening of the channel. Inward permeation (-Δμ) brings the inactivation gate back into the pore and allow the channel to switch back to a pressure-gated mode. Further pressure-mediated inward permeation leads the channel into a deep inactive state (inactivation gate tilted towards the center of the pore). Such transition is reversible and mediated further by outward permeation. (See also Figure S4)

To test our hypothesis we next stimulated patches with a pressure pulse at a positive voltage (+80mV) and again at the end of the mechanical stimulus we observed a sag followed by current reactivation (Figure 6B). The holding voltage was then stepped to +5 mV for 2 s to avoid inward permeation and to allow the channel to complete deactivate. If the stable conformational change after the sag phase corresponded to a transition between a pressure-gated and a voltage-gated mode it would be possible to re-open the channel by applying a simple depolarizing voltage protocol to mediate channel opening. We then stimulated the patches with a series of depolarizing steps (from 0-60 mV in 10 mV increments) without a mechanical stimulus. Strikingly, both the R2482H and R2482K mutant channels were directly activated by the voltage steps in a manner similar to classical voltage-gated ion channels with a V_50_ of 50.7 ± 9.3 (10 cells R2482H) and 19.5 ± 4.5 mV (5 cells R2482K) respectively (Figure 6 G). This experiment demonstrates that PIEZO1 carrying a human pathogenic mutation can transition from a pressure-gated mode to a voltage-gated mode provided that the inactivation gate is kept open. Repolarizing the patch back to negative potentials to allow inward ion permeation produced PIEZO channels that showed no voltage gating (Figure S4E). Under experimental conditions in which the inactivation gate remained open (no inward permeation after the mechanical stimulus) voltage-gated currents displayed a sigmoidal time course for activation and voltage dependency with a marked delay, especially with smaller voltage steps (Figure 6B, C). Thus PIEZO channels can be gated by either pressure or voltage. Voltage and pressure likely trigger channel opening via two different mechanisms as revealed by extrapolating the gating charges from the Boltzmann fits of the macroscopic conductance (Table S1). While in pressure-gated mode the equivalent gating charge for both R2482K and R2482H is approximately 2.2 e_0_, in voltage-gated mode the value reaches 6e_0_. This remarkable difference shows that in a voltage-gating mode a much higher number of charges must translocate across the membrane to drive channel opening.

Next, we asked whether wild type PIEZO1 channels can also transition from a pressure-gated to a voltage-gated mode. We found that less than <5% of the patches expressing the wild type PIEZO1 exhibited a reactivation which required high pressure and voltages as high as 180mV (Figure S4C). These extreme stimulation protocols often resulted in patch rupture. We thus chose to use a recently discovered small molecule modulator of PIEZO1 activity, Yoda1, to promote conformational states of the PIEZO1 that may more easily enter a voltage-gated mode. Yoda1 has been described as an efficient opener of PIEZO1 channels in artificial lipid bilayers but fails to directly gate the channel in whole cell configuration and it is a poor partial agonist in cell-attached patches (Syeda et al., 2015). In outside-out patches concentrations of Yoda1 close to the solubility limit of the compound (10μM) produced no appreciable channel opening when Yoda1 was applied either in the extracellular buffer or via the recording pipette (data not shown). However, as previously reported, Yoda1 acted as a gating modulator of PIEZO1 (Figure S4A,B) by slowing down both the current rise time, inactivation and deactivation time constants (Syeda et al., 2015). We therefore used 5μM Yoda1 in the recording pipette and asked whether under these conditions PIEZO1 could switch to a voltage-gated mode. Indeed PIEZO1 currents in the presence of Yoda1 showed a sag and a reactivation process starting at positive voltages (mean threshold was 43.3 ± 5.6 mV, 6 cells). We then performed a voltage protocol in absence of pressure as described for the R2482H and R2482K PIEZO1 mutants. Interestingly, under these experimental conditions we observed robust voltage activated outward currents with V_50_ of 31.9 ± 7.4 mV (5 cells). The speed of current activation increased exponentially with voltage (Figure 6 F) and an estimation of the gating charges from the steepness of the Boltzmann fit yielded a value of 5.2 e_0_.

We established that mammalian PIEZO channels can switch from mechanically gating mode to a voltage gated mode and that this transition is promoted by mutations near the pore region or by a synthetic gating modulator.

#### Evolutionary conservation of PIEZO voltage-gating

The *Piezo1* gene can be found in plants, invertebrates and protozoa (Coste et al., 2010). In insects a sensory role for *Drosophila melanogaster* PIEZO has been established as the channel is expressed in larval nociceptors and loss of function leads to reduction in mechanical nociception (Kim et al., 2012). The *Piezo1* gene identified in zebrafish (*Danio rerio)* has recently been found to be crucial for epithelial homeostasis (Eisenhoffer et al., 2012; Gudipaty et al., 2017). Here we cloned cDNAs encoding the Drosophila (DmPIEZO, accession number JQ425255), Zebrafish Piezo1 (ZPIEZO1) and the human PIEZO1 (hPIEZO1) in order to compare the voltage-dependent behavior of PIEZO channels across three orders of the animal kingdom. To study the orthologues under the same controlled experimental we the N2a *^Piezo1^*^-/-^ cells (see Methods and Figure S2). While the biophysical and mechanosensitive properties of the fly and human PIEZO1 have been extensively characterized (Kim et al., 2012; Zhao et al., 2016), this is the first report of the biophysical properties of heterologous expressed ZPIEZO1. ZPIEZO1 showed mechanosensitivity in outside-out patches with a pressure response behavior of P_50_ of 51.0 ± 2.7 mmHg (n = 9) which is typical for several PIEZO1 channels (Figure S5A). However, the inactivation time constant τ at -60mV and at saturating pressure was ~4 times slower than the mouse and human orthologues. Interestingly, in contrast to the rectifying behavior of mPIEZO1 (Figure 1A), the I/V relationship of ZPIEZO1 showed that this behaves as a perfectly ohmic channel at voltages between -100mV and 100mV. This suggests that profound structural differences in the ion permeation pathway exist between the ZPIEZO1 and the mPIEZO1.

As shown for mouse PIEZO1 the human channel (hPIEZO1) desensitized quickly in response to consecutive pressure stimuli (Figure 2A, Figure 7A). In contrast, the amplitude of DPIEZO and ZPIEZO1 currents only attenuated modestly with repeated pressure stimuli (Figure 7A). We next investigated voltage modulation of DmPIEZO and ZPIEZO1 using the tail current protocol (Figure 1D). The calculated apparent open probability of hPIEZO1 at <0 mV was very small (<5%) and thus almost identical to mPIEZO1 (Figure 7A). However, the apparent open probability of the channels from a pre-pulse holding potential of 0 mV differed dramatically between hPIEZO1, DmPIEZO and ZPIEZO1 (Figure 7C,D). The apparent open probability of DmPIEZO was only weakly modulated by voltage and we calculated that ~50% of the channels were available from a pre-pulse holding voltage negative to 0 mV (Fig. 7C,D). In contrast, the gating of ZPIEZO1 was completely unaffected by voltage so that 100% of the channels were available regardless of pre-pulse voltage. In this respect the virtual absence of voltage modulation of the ZPIEZO1 protein was very similar to the mouse PIEZO1 mutant R2482K (compare Figure 7D and Figure 4B). Considering the similarities in voltage modulation between the mouse PIEZO1 mutant R2482K and the fly and zebrafish PIEZO orthologues, we next asked if DmPIEZO and ZPIEZO1 could be directly opened by positive voltage steps. We applied the same combination of pressure and voltage protocols to patches from N2a *^piezo1^* ^-/-^ cells that expressed the orthologues as we had used for mouse PIEZO1. Mechanically-gated currents originated from the hPIEZO1 at voltages as high as 160mV caused the deactivation of hPIEZO1 to slow down but no sag or reactivation phase was observed (n=10 Figure S6). In contrast, mechanically-gated currents mediated by wild type DmPIEZO and ZPIEZO1 showed sag and reactivation kinetics (switch to voltage-gated mode) following pressure stimuli starting at 50mV and 20mV respectively (n=5, Figure S6). If the reactivation of the channels was indicative of a switch to voltage-gated mode then the protocol we previously used (step to 5 mV for 1-2 s) should allow voltage gating (Figure 6). Indeed, the opening of both DmPIEZO and ZPIEZO1 could be triggered by the application of voltage steps alone (Figure 7 E,F). Voltage-gating of the wild type ZPIEZO1 protein was robust and very similar to wild type PIEZO1 in the presence of the synthetic gating modulator Yoda1 (Figure 6), and showed a V_50_ of 20.2 ± 6.8 mV (n=7) and a slope of 7.9 ± 1.3 with a corresponding gating charge of 3.8 ± 0.7. The DmPIEZO protein was also opened by positive voltage pulses, but showed markedly slower activation kinetics and voltages as high as 140mv still did not produce saturating activation (Figure 7E,F). Consequently, the voltage sensitivity of DmPIEZO was shifted very significantly to the right compared to ZPIEZO1. Thus, PIEZO channels in both Drosophila and zebrafish can be switched to a voltage-gated mode even more readily than mammalian PIEZO proteins.

**Figure 7.**
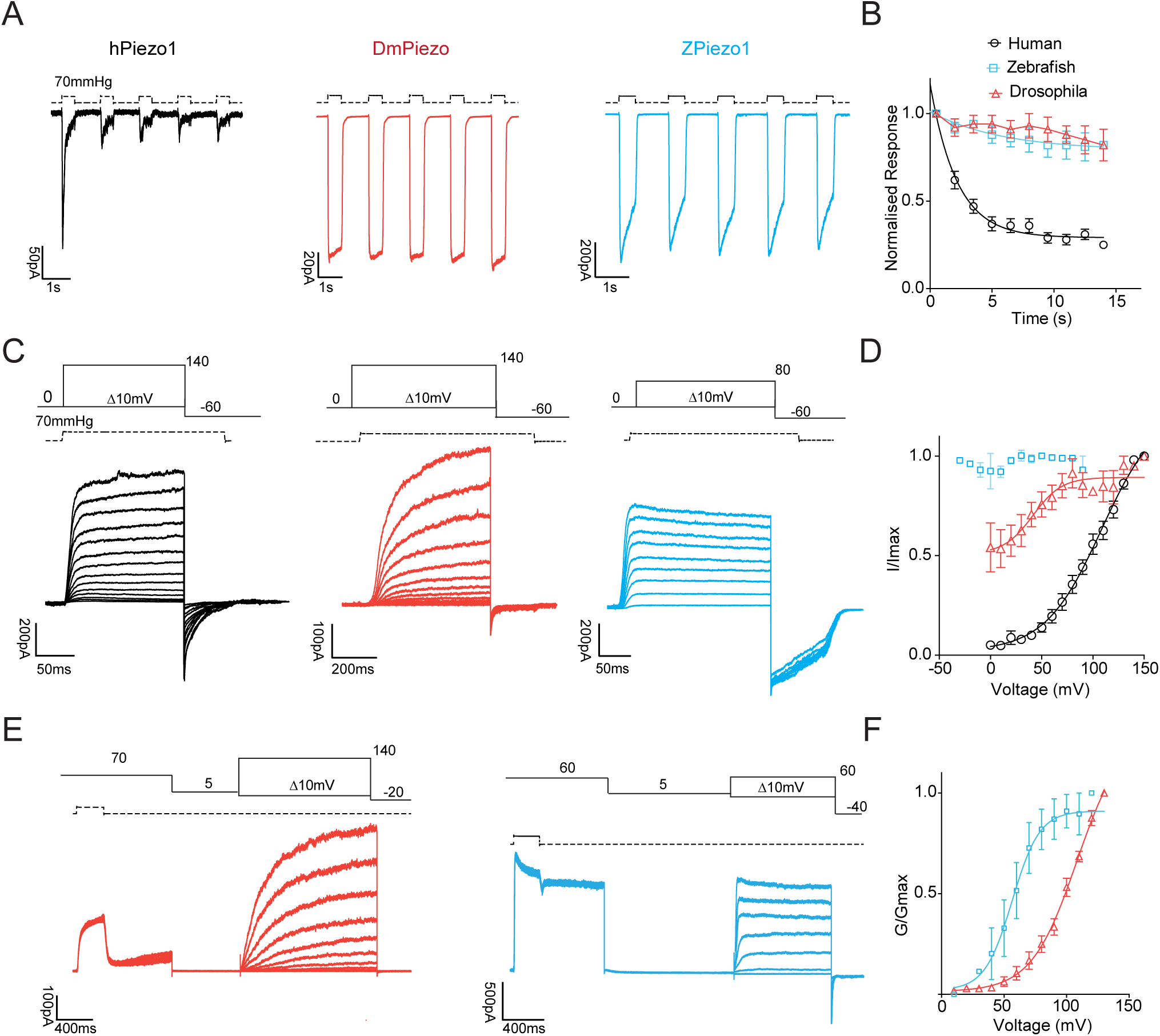
Properties of human, drosophila and zebrafish PIEZO1. A) 5 pulses of saturating pressures at -60mV were applied to patches pulled from N2a *^Piezo1^*^-/-^ expressing the human (black), drosophila (red) and zebrafish PIEZO1 (blue). B) Current amplitudes were normalized to the first pressure pulse and plotted against time. C) Current responses to a pressure stimulation of 70mmHg during 300ms voltage steps ranging from 0 to 140mV, followed by a repolarization step to -60mV to obtain tail currents. D) Tail currents from individual cells were normalized to their maximum and fitted to the Boltzmann relationship (human V_50_ = 96.7 ± 4.8 mV, slope 19.9 ± 0.9, 5 cells, drosophila V_50_ = 40.0 ± 9.8 mV, slope 12.1 ± 3.9, 7 cells). Note how the pressure mediated currents of the zebrafish PIEZO1 are insensitive to voltage. Pooled data are shown as mean ± SEM. E) Same stimulation protocol as in Fig 6B for DPIEZO (red) and ZPIEZO1 (blue). F) Macroscopic conductance values were normalized to maximum conductance and plotted against voltage to obtain a G/V curve. DPIEZO did not reach saturation at voltages as high as 140mV, while ZPIEZO had V_50_ of 20.2 ± 6.8 and a slope of 7.9 ± 1.3 (n = 7). (See also Figure S2, S5 and S6)

## DISCUSSION

Here we have shown that both vertebrate and invertebrate mechanosensitive PIEZO ion channels are polymodal, being voltage modulated and under circumstances (causing pathological conditions in humans) even gated directly by voltage in the absence of mechanical stimuli. One important new feature we have identified in mammalian PIEZOs is that under physiological resting membrane potentials ~95% of the channels are unavailable for pressure gating. However, outward permeation of the channel driven by depolarizing voltage rapidly removes both desensitization and the loss of inactivation that is observed with repetitive mechanical stimulation. Indeed, with outward permeation driven by stronger electrochemical gradients almost all the PIEZO channels can recover their mechanosensitivity. The voltage modulation and voltage gating of PIEZO channels appears to be a fundamental property of the channel pore and is dramatically changed by well characterized pathological mutations that cause Xerocytosis. We have also shown that pressure stimuli given at positive potentials can transition the channel into a voltage gated mode, a situation strongly promoted by pathogenic mutations and by a synthetic gating modulator, Yoda1 (Syeda et al., 2015). Our data can be explained by the existence of an inactivation gate for which a closed conformation is strongly favored at physiological resting membrane potentials. The open conformation of the inactivation gate is on the other hand promoted by outward permeation of the pore, and in this conformation direct voltage gating of the channel is possible. Direct voltage gating is a very prominent physiological feature of older vertebrate (fish) and invertebrate (fly) PIEZO proteins. The susceptibility of PIEZO channels to voltage modulation or voltage gating are thus ancient properties of the channel that differ in prominence throughout the evolution of the channel family. Voltage modulation and voltage gating of PIEZO channels is likely a deep property of such channels that has been adapted to enable a variety of mechanosensing roles in many cell types.

A prominent feature of PIEZO1 currents is that they display desensitization and a loss of inactivation with repeated stimuli (Coste et al., 2010; Gottlieb et al., 2012). This property could be a problem for mechanosensing under circumstances where mechanical force should be constantly monitored over time. We now show that slowing of inactivation and desensitization can both be rapidly reversed by outward permeation of the channel which may in fact happen in excitable cells under physiological conditions allowing them to re-acquire mechanosensitivity. Our data can be explained by the presence of an inactivation gate that, similarly to that shown in K2P channels (Schewe et al., 2016), is closed by inward ion flux and re-opened by outward permeation. We also show that outward ion permeation allows PIEZO1 to exit desensitization through a slow transition (time course of 100 ms, at least one intermediate state) (Figure 2E).

By using conditioning positive voltage steps we found that it requires considerable driving force to open the inactivation gate, > +80mV to fully activate all channels (Figure 3A,B). Nevertheless, mechanical activation of PIEZO channels combined with outward permeation shifts more channels into a non-desensitized state available to sense mechanical force. Thus voltage modulation of PIEZO channels is potentially a powerful mechanism to regulate force sensing in cells, especially considering that in the absence of outward permeation of the channel ~ 95% of PIEZO channels cannot be opened by mechanical force (Figure 2). Indeed it has been shown that mechanosensitive currents in sensory neurons can be sensitized by inflammatory signals (Lechner and Lewin, 2009; Lewin et al., 2014) and this could take place via regulation of PIEZO channels (Dubin et al., 2012). It is thus conceivable that inflammatory signals could shift voltage sensitivity of the PIEZO channels into a range where action potential firing in the presence of mechanical force could release many more channels for activation by opening the inactivation gate.

Despite the high amino acid sequence homology between PIEZO1 and PIEZO2 these two channels appear to be activated by different kinds of mechanical stimuli. Whole cell patch clamp experiments in cells has provided evidence that expression of PIEZO1 or PIEZO2 is accompanied by a fast current activated by cell indentation or substrate deflection (Coste et al., 2010; Poole et al., 2014). PIEZO1 channel activity has also been extensively studied in excised patches and in artificial lipid bilayers where, similarly to TRAAK channels (Brohawn et al., 2014), membrane stretch or increased curvature are sufficient to gate the channel (Syeda et al., 2016). So far evidence that PIEZO2 channels can be gated by membrane stretch has been lacking. Furthermore, attempts to elicit pressure-mediated currents in outside out patches from cells overexpressing PIEZO2 failed both in a heterologous system (this report) and in Merkel cells, where PIEZO2 is abundantly expressed (Ikeda and Gu, 2014; Woo et al., 2014). Here we show using a PIEZO1/PIEZO2 chimeric expression construct that the N-terminal region of PIEZO1 allowed us to measure pressure activated currents in excised patches from N2A cells in which the large portions of *Piezo1* gene had been deleted (Figure S2). Importantly, the kinetic properties of the mechanosensitive current were characteristic of PIEZO2 currents displaying faster inactivation kinetics than PIEZO1 (Figure 5, Figure S3C). Interestingly, the pore region of the chimeric P1/P2 protein was capable of sensing voltage and outward permeation could also relieve loss of inactivation and desensitization. Indeed, we also found that at resting membrane potentials (-60mV) the apparent open probability of the chimeric protein was very low (~5-15%) similar to that of PIEZO1. Thus the last 350 amino acids of PIEZO2 appear to code for voltage modulated pore with a putative inactivation gate that can be activated by membrane stretch when fused with N-terminal PIEZO1 sequences. These experiments support the view that force may be transferred to the pore forming C-terminal regions via sequences in the N-terminal region (Zhao et al., 2017), but in contrast to other reports our experiments are not complicated by the presence of PIEZO1 (Zhao et al., 2016).

Xerocytosis is a human genetic disease that has been associated with missense mutations in PIEZO1. Despite extensive *in vitro* studies the only effect that has been described is a gain of function in the inactivation kinetics. Here we have shown that the strongest Xerocytosis mutants have large effects on the voltage sensitivity of the channel, thus the R2482H/K mutation shifts the voltage modulation 60 mV leftwards to dramatically increase apparent open probability at negative potentials (Figure 4). The mutated Arginine residue is located at the bottom of the inner helix of PIEZO1 a position close to the inner pore. Interestingly, in other ion channels families Arginine residues at the extremities of pore lining regions have been shown to be essential for anchoring the helices to the lipid bilayer and to stabilize the closed conformation of the channel (Moorhouse et al., 2002). We thus speculate that R2482 normally stabilizes the channel in a closed conformation and movement of this residue (together with other basic ones) might confer voltage-modulation. It is worth noting in this context that Arginine residues in the S1-S4 domains play a fundamental role in voltage sensing in most canonical voltage-gated ion channels (Arhem, 2004; Liman et al., 1991; Swartz, 2008). Interestingly, voltage modulation of pressure-gated ion channels has previously been described for the archetypical pressure sensitive bacterial channel MscS, which also requires an arginine residue within the lipid bilayer (Bass et al., 2002; Jiang et al., 2003; Martinac et al., 1987). However, so far there are no reports that bacterial mechanosensitive channels can be directly voltage-gated.

Our data show that PIEZO1 is a polymodal channel that can be gated directly by voltage. A mutant channel involved in human disease (mouse R2482H/K) undergoes an unprecedented switch from a pressure-gated to a voltage-gated mode at voltages more positive than ~30mV. The transition is caused by a sustained outward ion flux in the absence of pressure. The switch between the two modes is characterized by a sag followed by a reactivation phase whose occurrence is purely voltage dependent (Figure S2D). We hypothesize that switching may cause the inactivation gate to remain fully open and that PIEZO1 undergoes a conformational change that allows the channel to directly respond to voltage. Evidence for a profound conformational change was provided by estimating the gating charge (e_0_) as to sense voltage charged residues position within the electric field change upon voltage sensor activation. We estimated the gating charges from the slope factor of macroscopic currents to provide a lower limit of the charge movement. However, we observed a 3-fold increase in e_0_ upon switching from pressure to voltage gating mode (from 2.2 to ~6 e_0_ for R2482K, Supplementary Table 1), suggesting that charges not available to move with electric field changes in pressure mode become free to move in voltage gated mode because of a large conformational change near the pore. A graphical explanation of this model is given in Figure 6H. We have shown that voltage gating and voltage modulation of PIEZO channels is an ancient property present in both invertebrate (fly) and older vertebrate (fish) wild type PIEZO channels. Comparisons of the pore regions of these channels could provide profound insights into the structural determinants necessary for the inactivation gate and its regulation by outward permeation. In summary, the revelation that voltage can modulate and gate mechanosensitive PIEZO channels is likely a deep property of this channel family that is integral to their physiological function across a wide variety of mechanosensing cells.

## Author Contributions

Conceptualization MM, GRL. Methodology MM. Investigation, MM, MRSV,RF. Writing MM,GL. Visualization MM, RF. Supervision MM, GRL. Project Administration MM. Funding Acquisition GRL.

## Acknowledgments

We thank Prof Thomas Baukrowitz for initial comments on the manuscript. Liana Kozitzki for technical assistance.

## STAR Methods

### Cell culture

The mechanosensitive cell line, N2A, was cultured in DMEM/Opti-MEM (50/50) containing 5% fetal calf serum (FCS) and 1 % penicillin and streptomycin. The medium was changed to a serum free medium before transfection. Cells were transfected with mouse Piezo1 or Piezo2 cDNA constructs containing an IRES-EGFP sequence to select transfected cells (a gift from Dr. Ardem Patapoutian). Cells were transfected in a 35mm petri dish, pre-coated with poly-(L)-Lysine, with 1μg DNA and 3μl of HD Fugene (Promega) following the manufacturer’s protocol and were recorded 12 to 48 hours after transfection.

### Molecular Biology

R2482H and R2482K were introduced in the mouse Piezo1 plasmid using the XL QuikChange^TM^ Site Directed Mutagenesis (Agilent Technologies). The chimera between Piezo1 and Piezo2 was produced by PCR by joining residues 1-2188 of mPiezo1 and 2472-2822 of mPiezo2. The resulting chimeric construct of 2539 amino acids was cloned into an IRES-EGFP containing vector by SLIC reaction.

The *Drosophila melanogaster* Piezo was cloned from Drosophila mRNA (a gift from Dr Robert Zinzen), while the *Danio rerio* (zebrafish) Piezo1 cloned from individuals at 48 hours post fertilization (a gift from Dr Daniela Panakova).

### Generation of Piezo1 KO N2a cells

Deletion of the *Piezo1* gene was generated using CRISPR/Cas9 technology. 4 gRNAs targeting exon 6 and exon 45 were generated using the gRNA designer (http://crispr.mit.edu/) for the nickase Cas9 (Addgene plasmid 48140, a gift from Feng Zhang). N2a cells were transfected with gRNA and Cas9n and single cell sorted 1 week after transfection to generate a clonal *Piezo1* -/- cell line. The selected clone had a deletion of 17 Kb and no mechanically activated currents were detected by using soma indentation (Figure S2). Sequences of the gRNAs and the sequencing reactions of the resulting deleted alleles are reported in Figure S2.

### Electrophysiology

Recordings were performed in excised outside-out patches pulled from N2A cells. Experiments were performed at room temperature. Recording pipettes were prepared using a DMZ puller and subsequently polished to a final resistance of 6-8 MΩ. Pressure stimuli were applied through the recording pipette via a High Speed Pressure Clamp (Ala Scientific). Soma indentation experiments were performed in whole-cell configuration using microelectrodes with a resistance between 2 and 4 MΩ. Uncompensated series resistance values were less than 2 MΩ. Cells were held at -60mV and mechanical stimulation was performed using a blunt glass probe (tip size 3-4μm).

Experiments were performed in symmetrical ionic conditions and in a divalent-free buffer. Solutions contained (in mM): 140 NaCl, 10 HEPES, 5mM EGTA adjusted to pH 7.4 with the NaOH. Currents were sampled at 10 kHz and filtered at 3 kHz using an EPC-10 amplifier and Patchmaster software (HEKA, Elektronik GmbH, Germany). The currents were subsequently analyzed using FitMaster (HEKA, Elektronik GmbH, Germany). To calculate the single channel conductance of PIEZO1, stretch of openings were first filtered at 1 kHz, subsequently all point histograms were constructed using FitMaster. From a double Gaussian fit the single channel conductance was calculated by plotting the average current at each potential versus the voltage. Slope conductance values at positive and negative holding potentials were obtained by plotting current-voltage relationships, and by fitting the following relationship to the data points (each point was the average of 50 to 200 openings)

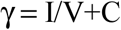

where γ is the slope of the line, I is the single current measured, V is the voltage and C is the reversal potential (Prism 5, Graphpad).

Saturating tail current amplitudes were fitted to a standard Boltzmann equation:

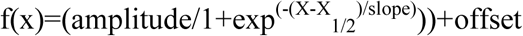

where the slope corresponds to RT/zF (R, universal gas constant; T temperature; z equivalent gating charge; F, Faraday constant).

### Statistics

All data sets were tested for normality: parametric data sets were compared using a two-tailed, Student’s *t*-test, paired or unpaired depending on the experimental setup, nonparametric data sets were compared using a Mann-Whitney test. One-way ANOVA and Dunnett’s *post-hoc* test were used as indicated in the figure legends.

**Figure S1.**
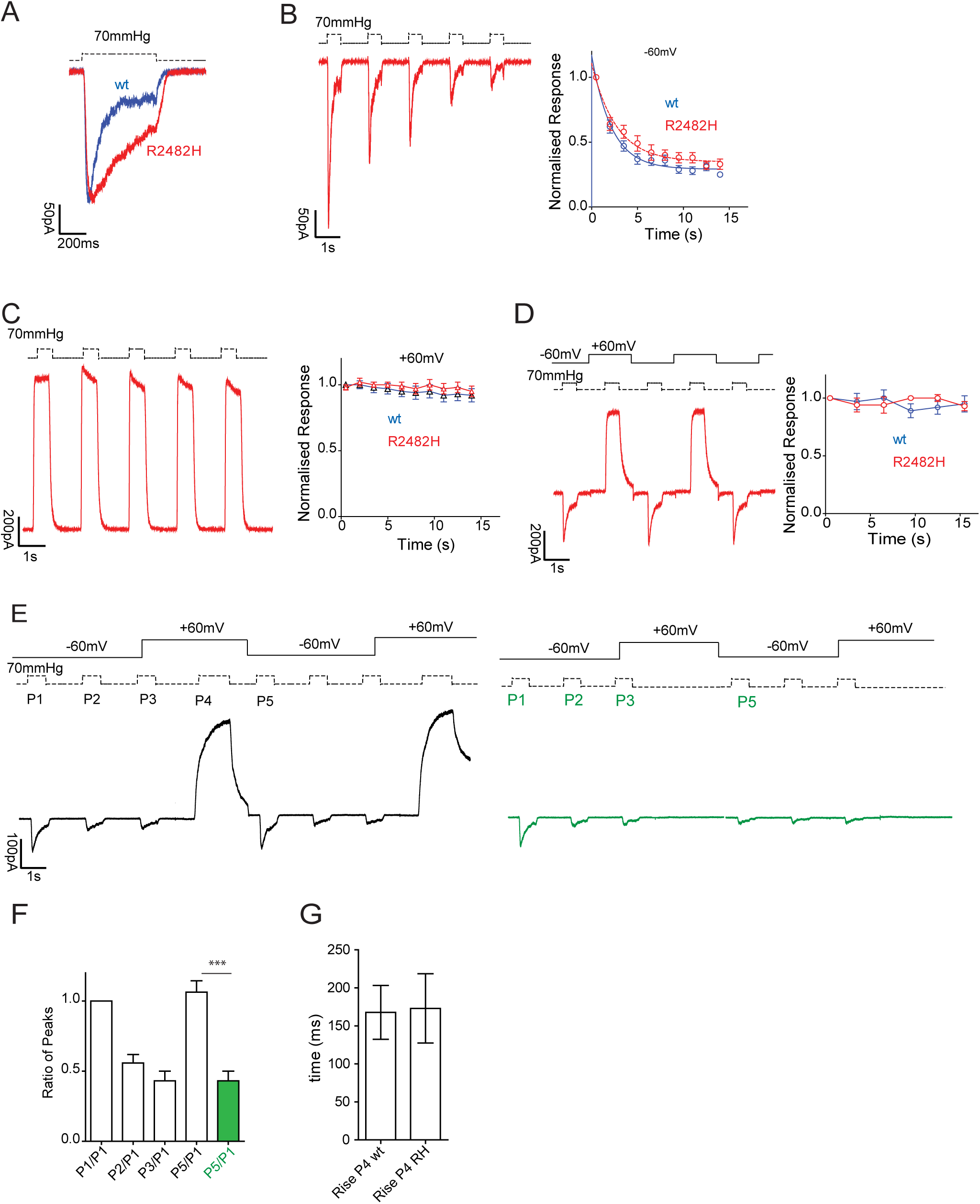

**Figure S2.**
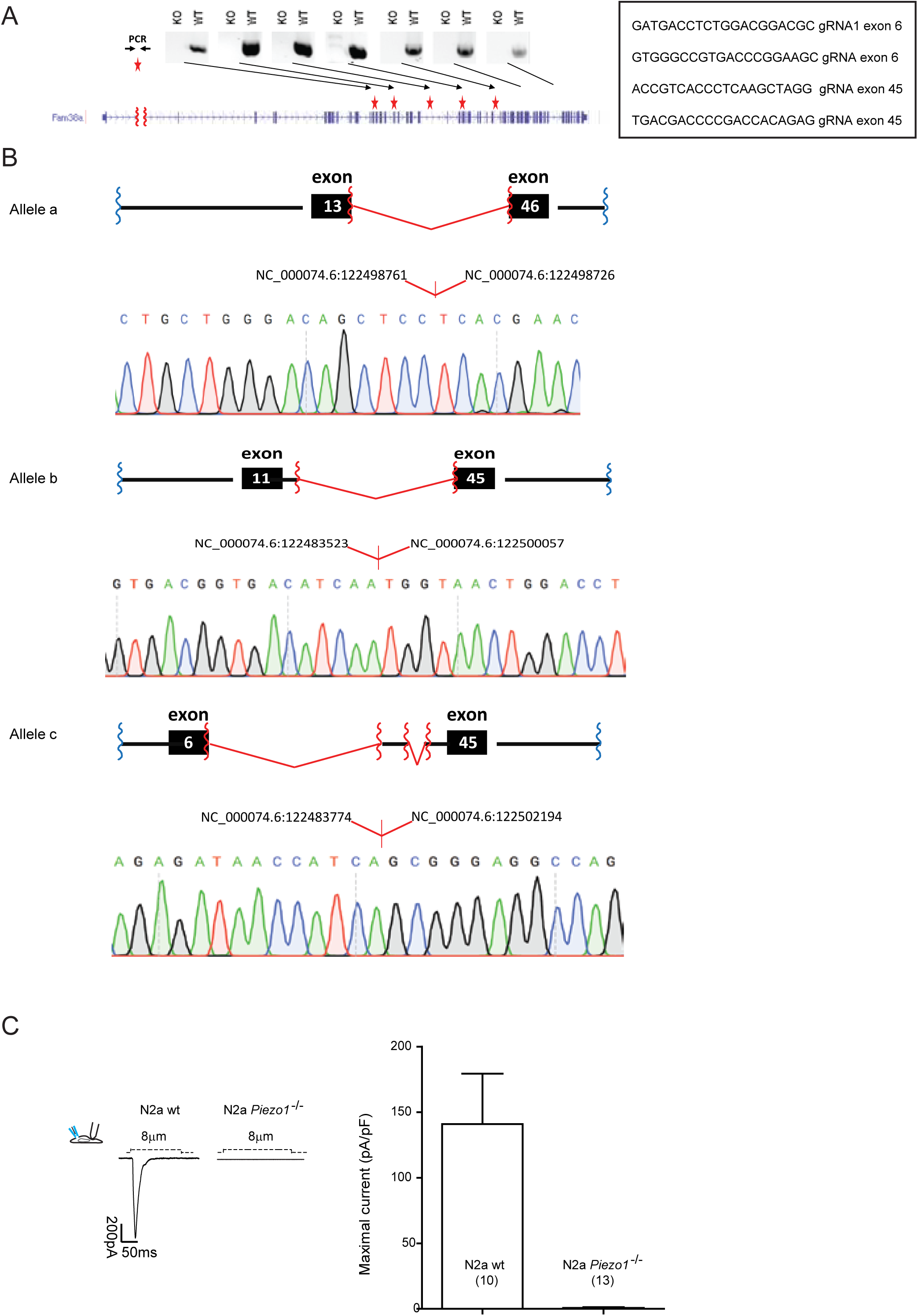

**Figure S3.**
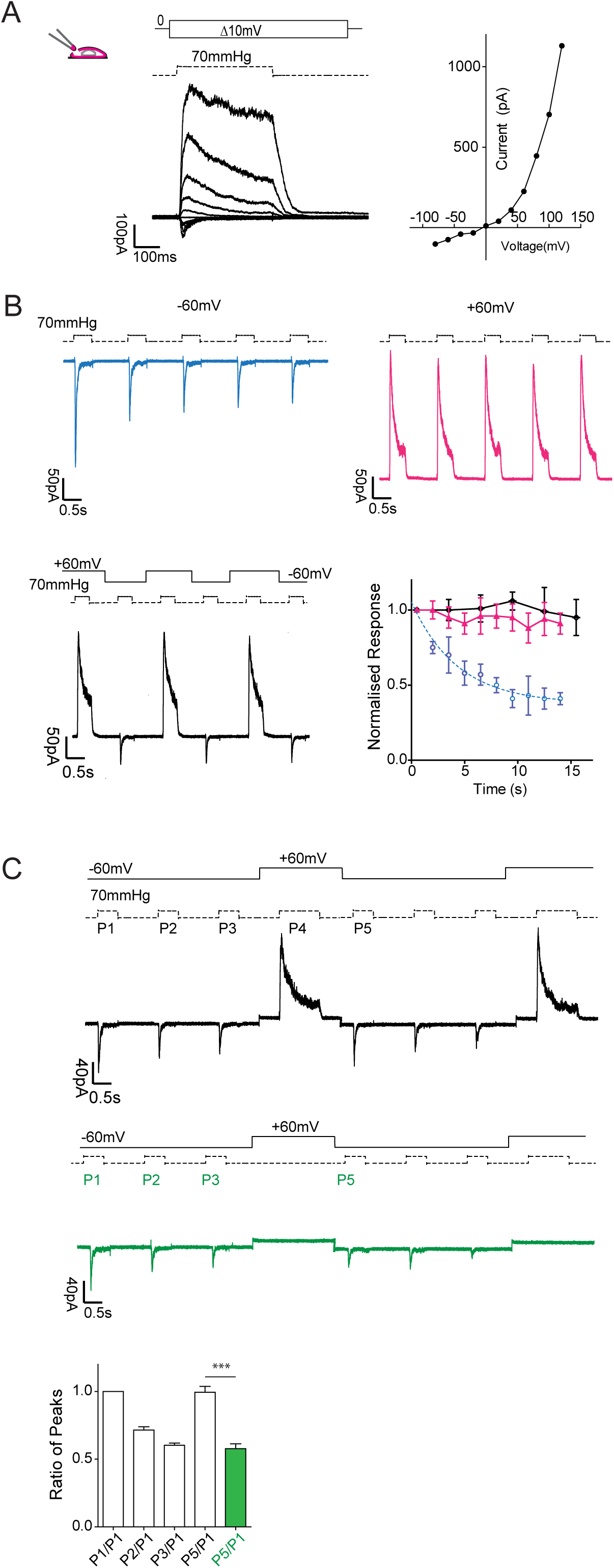

**Figure S4.**
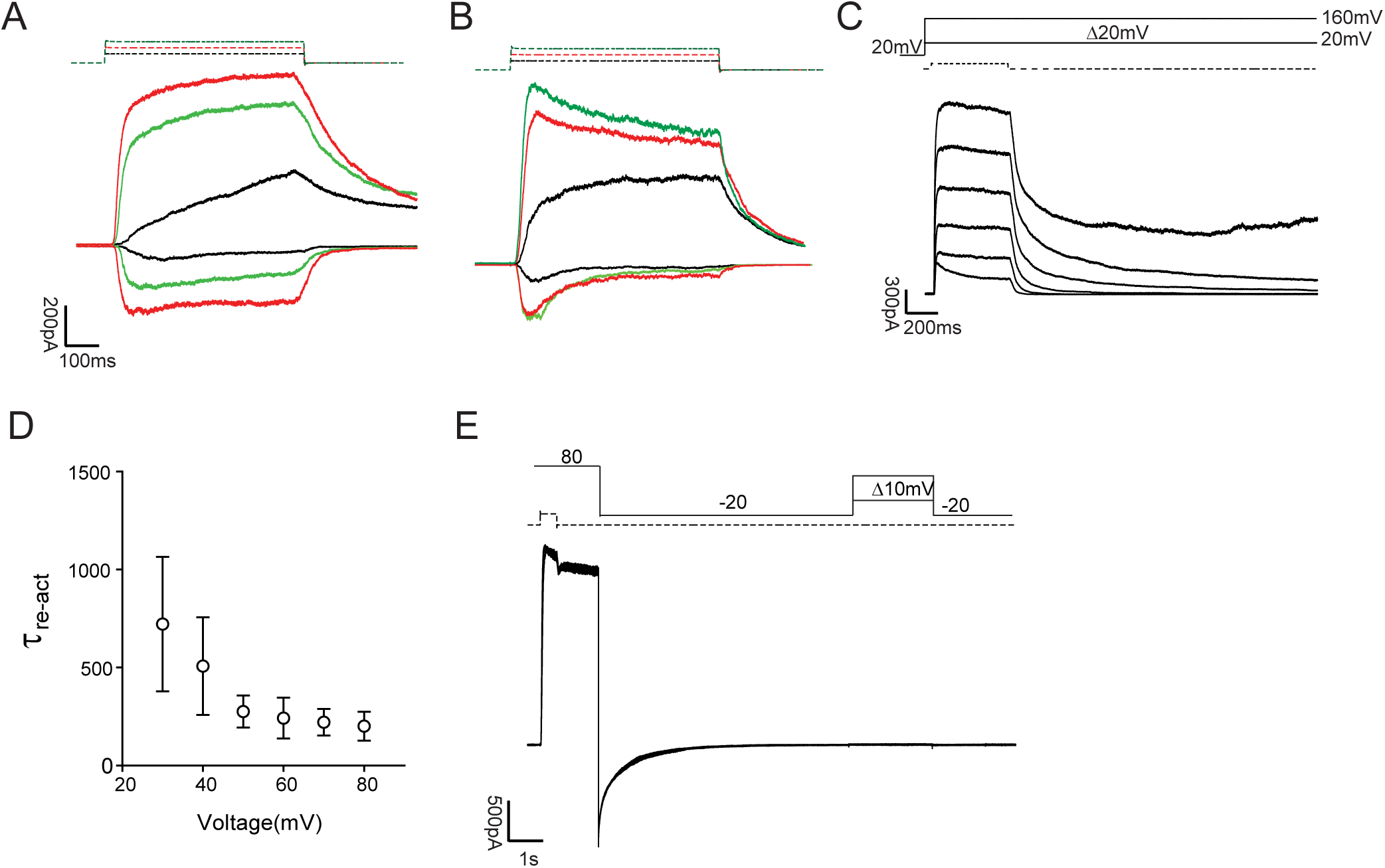

**Figure S5.**
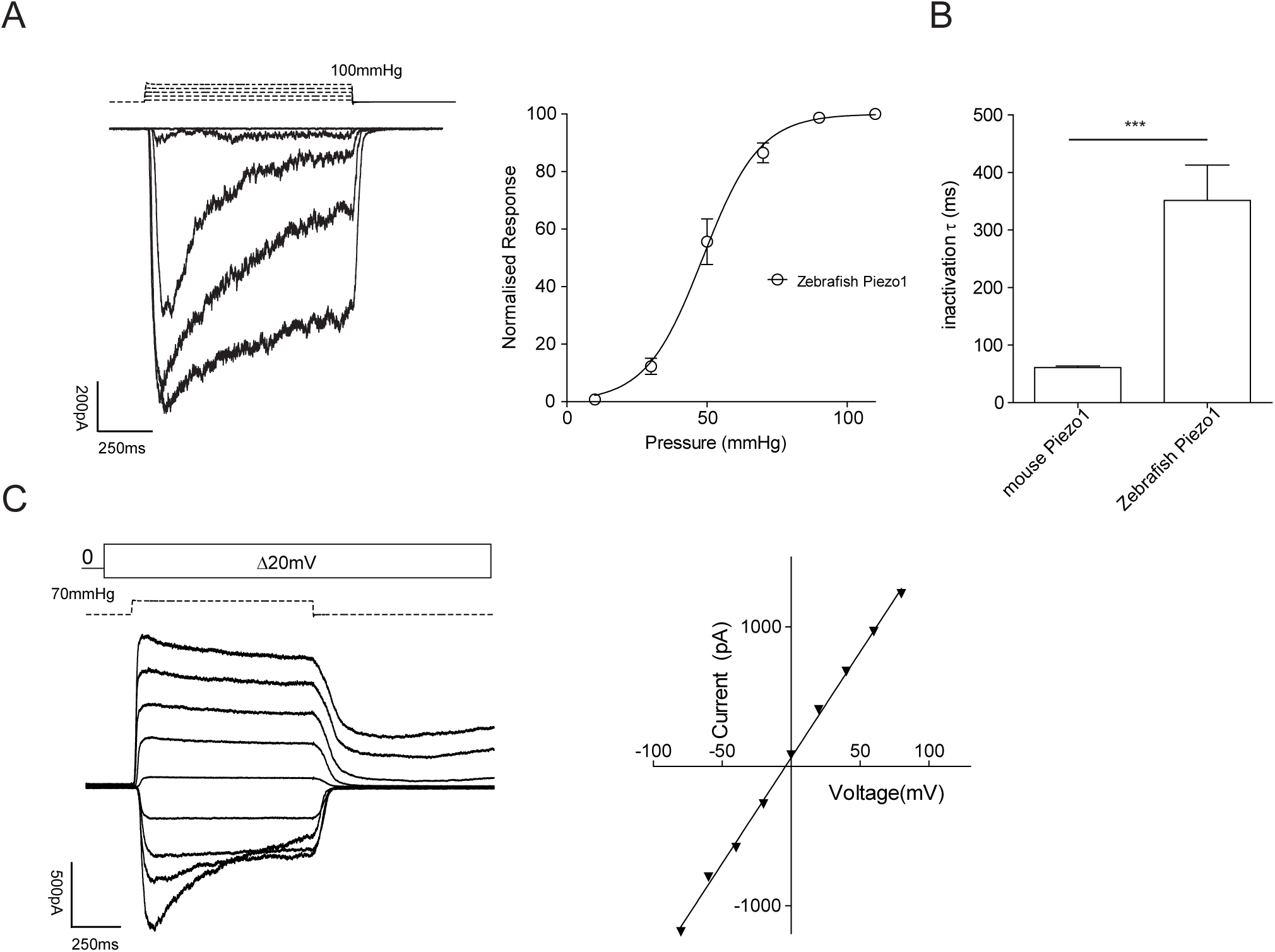

**Figure S6.**
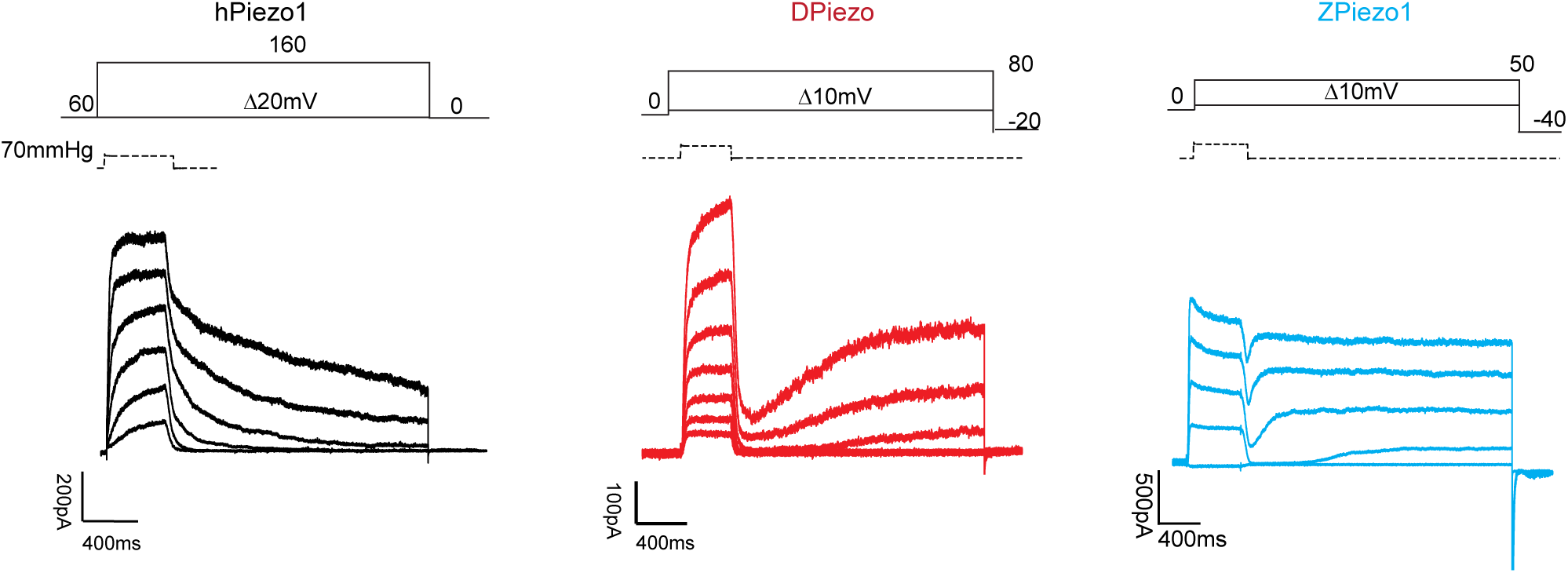

